# Analysis of persistence thresholds for a nonlocal PDE–ODE model of bacterial persister cells

**DOI:** 10.64898/2026.04.20.719571

**Authors:** Chongming Li, Tyler Meadows, Troy Day

## Abstract

Within many bacterial colonies, persister cells exist as a subpopulation that is tolerant to antibiotics and other stressors, yet not genetically distinct from the rest of the colony. A recent study has proposed epigenetic inheritance as a mechanism that leads to the presence of persister cells. We analyze a nonlocal PDE–ODE model introduced in that study to describe the epigenetic inheritance process and establish its mathematical well-posedness, including existence, uniqueness, and nonnegativity of solutions. We identify a sharp parameter threshold delineating extinction from persistence of the colony: below this threshold the washout equilibrium is globally asymptotically stable, while above it a unique positive equilibrium exists and the population is weakly persistent. Notably, this threshold is independent of the internal community structure.

## 1. Introduction

Bacterial persister cells are a subpopulation of some bacterial colonies, exhibiting resistance to certain stressors such as antibiotics and starvation. Persister cells were first discovered in 1944 by Joseph Bigger [3] during his experiments with penicillin. After treating a colony of bacteria with the drug, a small number of cells would survive. New colonies grown using the surviving cells were just as susceptible to antibiotics as the original colony.

Persister cells survive the application of antibiotics and other stressors by entering a dormant, or quiescent, state. Since most *β*-lactam antibiotics, like penicillin, act by disrupting cell wall formation during cell division, avoiding replication entirely is an effective strategy for surviving antibiotics. Similarly, shutting down most cellular functions serves as an effective survival strategy during times of starvation, oxidative stress, and other environmental stresses.

Biological studies [8, 10, 16, 18] have shown that bacterial persistence is a phenotypic state rather than genetic adaptation; persister cells are genetically identical to the original wild-type bacteria, but differ in certain quantitative traits that enhance survival under stress. In mathematical models, these phenotypic traits are often represented by expression levels of particular chemicals, and it is assumed that bacteria switch to the persister state when the expression level exceeds a certain threshold [4].

Modelling persister cells via PDEs in the bacterial literature is relatively new as the existing models consider the interaction between persister and susceptible states of bacteria using ODEs (see [2]). The advantage of PDE models is the feasibility of tracking the exchange between subpopulations indexed by a continuum of expression levels, together with the dormant persister cells, at any given time. Furthermore, Day in [4] incorporated the birth-jump process via a nonlocal term. In spatial ecology, the term birth-jump process [7] is often used to describe situations in which newly produced offspring disperse from their original location immediately after birth. In our model, the change in phenotypes between generations can be interpreted, by analogy, as a birth-jump process.

We consider a model introduced in [4] consisting of an ODE coupled to a PDE with a nonlocal birth-jump term:

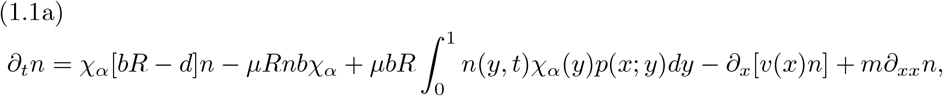

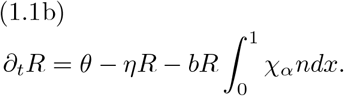

In system (1.1), we denote by *n* := *n*(*x, t*) and *R* := *R*(*t*), respectively, the density of individuals with expression level *x* ∈ Ω = (0, 1) (normalized to the unit interval) and the amount of resource present at time *t* ∈ ℝ_≥0_.

The first term of (1.1a), *χ*_*α*_(*x*)[*bR*(*t*) − *d*], represents the net per capita reproduction rate. We suppose that the birth rate is proportional to available resource *R*(*t*), while the death rate is a fixed constant *d*. The function

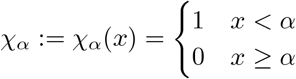

delineates between regular cells and persister cells; cells with expression levels above *α* are dormant and so do not replicate or die, while cells with expression levels below *α* behave normally.

The second and third terms of (1.1a) are related to changes in expression levels across generations during the reproduction process. By the biological interpretation in [4, 15, 17], the expression level can be transferred from parents to offspring during reproduction. We denote by *µ* the fixed probability of a change in expression level when a parent gives birth to an offspring. Inside the integral, *p*(*x*; *y*) is the probability density of the offspring’s expression level being *x*, given that the parent has expression level *y*, and assuming that a change takes place during the process of reproduction.

The final two terms of (1.1a) are advection and diffusion terms describing within-generation change. The advection term is based on a mechanism of cellular homeostasis [9] while the diffusion represents within-generation noise. Cellular homeostasis arises from microbes regulating their internal environment through physiological processes. The expression level of an individual is typically centered around a value that supports replication. This behavior is captured by choosing *v*(*x*) to model within-generation adjustment.

For the resource equation (1.1b), the parameter *θ* is a resource inflow rate whereas *η* is the per capita loss rate of the resource. The integral 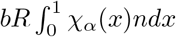 is the amount of resource consumed by the whole population of microbes per unit time.

Given the above biological description, we make the following assumptions: 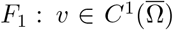 with *v*(0) = *v*(1) = 0, and there exists a unique point *γ* ∈ (0, 1) with *v*(*γ*) = 0, called the *homeostatic point*. A concrete example is the cubic *v*(*x*) = *v*_0_ *x*(*x* − *γ*)(*x* − 1) with *v*_0_ ∈ ℝ.

*F*_2_ : *p* ∈ *L*^2^(Ω × Ω) with *p* ≥ 0 a.e. and 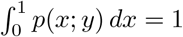 for a.e. *y* ∈ Ω.

*F*_3_ : *b, d, θ, η, m >* 0 and *µ* ∈ [0, 1].

Assumptions *F*_1_–*F*_3_ are in force throughout the paper, so we shall not repeat them in the statements of our results.

Lastly, we assume the following boundary and initial conditions:

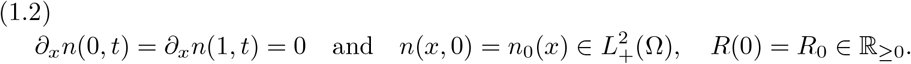

The main objective of this paper is to conduct a rigorous mathematical analysis of the nonlocal PDE–ODE system (1.1)–(1.2). The model is nonlinear, includes a discontinuous coefficient *χ*_*α*_, a nonlocal birth-jump operator, and an advection term with degenerate coefficients at the boundary. The lack of regularity in *χ*_*α*_ and the presence of a nonlocal operator complicate the analysis and preclude classical solutions around *x* = *α*. For this reason, we formulate the PDE component of (1.1) in the framework of *mild solutions* and analytic semigroup theory. This approach allows us to treat the linear diffusion-advection-reaction operator as the generator of a positive analytic semigroup and to incorporate the nonlinear and nonlocal terms via the variation of constants formula.

Our main results are as follows. We prove that system (1.1)–(1.2) is globally well-posed for nonnegative initial data, with solutions remaining nonnegative for all time (Theorem 2.7). The asymptotic behavior is governed by a sharp threshold: when *θ/η < d/b*, the washout equilibrium (0, *θ/η*) is the unique nonnegative steady state and is globally asymptotically stable (Theorem 3.9); when *θ/η > d/b*, a unique positive equilibrium exists (Theorem 3.10) and the population is weakly persistent (Theorem 3.12). Notably, this threshold depends only on the resource and demographic parameters *θ, η, d, b* and is independent of the advection velocity *v*, the redistribution kernel *p*, the switching probability *µ*, the persister cutoff *α*, and the diffusion coefficient *m*.

The paper is organized as follows: in Section 2, we investigate the well-posedness of system (1.1) by reframing the equations in a way that permits the use of semigroup theory [5, 12]. In Section 3 we identify conditions for the existence of steady state solutions to (1.1) and analyze the long-term behavior of solutions. In particular, we show that for values of 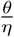 large enough, the system is weakly persistent; the total microbial biomass does not decay to zero as time tends to infinity. Finally, in Section 4, we illustrate our findings with numerical simulations, which help support the conjecture that the positive steady state is locally asymptotically stable when the system is weakly persistent.

## 2. Well-Posedness of the Model

Since *χ*_*α*_(*x*) is discontinuous, we work in an *L*^2^(Ω) framework and formulate solutions in the mild sense using semigroup theory. For *T >* 0, define the Banach spaces

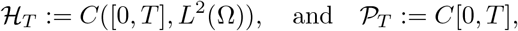

with norms

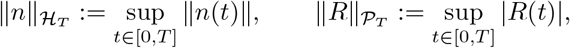

respectively. Throughout, ∥ · ∥ and ⟨·,·⟩ denote respectively the norm and inner product on *L*^2^(Ω); other norms carry explicit subscripts. For fixed *R*(*t*) ∈ 𝒫_*T*_, equation (1.1a) can be seen as a non-autonomous, nonlocal parabolic PDE. The right hand side of (1.1a) may be split into an autonomous part,

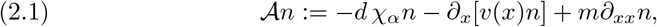

with domain

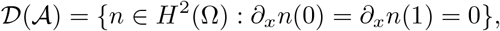

and a non-autonomous part *bR*(*t*)𝒩 *n*, where

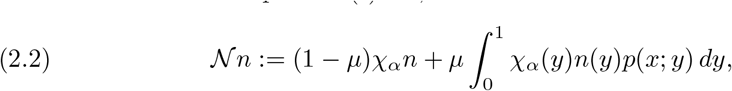

with domain 𝒟(𝒩) = *L*^2^(Ω).

We first study some preliminary semigroup properties of the operator 𝒜.

### Lemma 2.1.

*The operator* 𝒜 *given in* (2.1) *generates a quasi-contraction semi-group. In particular, there is a constant ω* ∈ ℝ *such that*

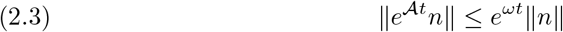

*for all t >* 0 *and all n* ∈ *L*^2^(Ω).

*Proof*. We complete the proof by verifying the conditions of [6, Theorem 9.5.2]. The domain 𝒟 (𝒜) is dense in *L*^2^(Ω) because 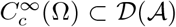 (compactly supported smooth functions trivially satisfy the Neumann conditions) and 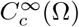 is dense in *L*^2^(Ω). The closedness of 𝒜 will follow from the Lumer–Phillips conditions verified below [5, Theorem II.3.15]. For quasi-dissipativity, we must show Re ⟨*n*, 𝒜*n* ⟩ ≤ *ω* ∥*n*∥ ^2^ for *n* ∈ 𝒟 (𝒜). Consider the bilinear form induced by −𝒜 on *V* = *H*^1^(Ω). Using integration by parts and the assumption *v*(0) = *v*(1) = 0:

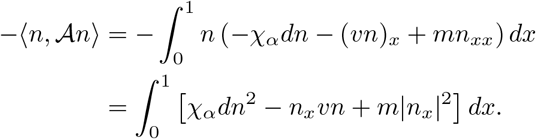

Using Young’s inequality 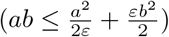 on the advection term 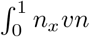:

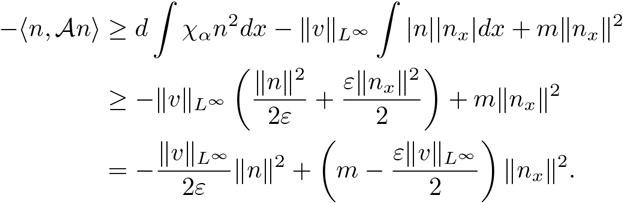

Letting 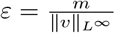, the coefficient of ∥*n*_*x*_∥ ^2^ becomes *m/*2. Thus:

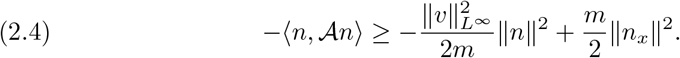

Rearranging gives 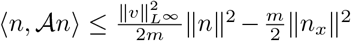. Defining 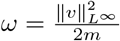, we have Re⟨*n*, 𝒜*n*⟩ ≤ *ω*∥*n*∥^2^. Next, we verify the resolvent condition. We show that (*λ* −𝒜) is surjective for *λ > ω*. Consider the equation *λn* − 𝒜*n* = *f* for *f* ∈ *L*^2^(Ω). The associated bilinear form *B* : *H*^1^ × *H*^1^ → ℝ is:

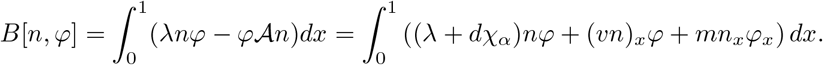

Using the preceding estimate (2.4), for any *n* ∈ *H*^1^(Ω):

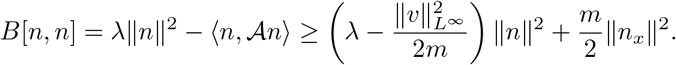

For 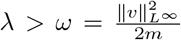, there exists *C >* 0 such that 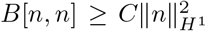. Thus, *B* is coercive. Since *B* is also bounded, by Lax-Milgram Theorem [14, Theorem 6.42], there exists a unique weak solution *n* ∈ *H*^1^(Ω). By standard elliptic regularity theory (see [14, Theorem 8.30]), since *v* ∈ *C*^1^ and *m >* 0, this weak solution satisfies *n* ∈ *H*^2^(Ω) and the boundary conditions. Thus, *n* ∈ 𝒟 (𝒜), and the range of (*λ* − 𝒜) is the entire space *L*^2^(Ω). In conclusion, conditions 1, 2, and 3 of [6, Theorem 9.5.2] with *H* = *L*^2^(Ω) are satisfied. Therefore, 𝒜 generates a quasi-contraction semigroup.

The following positivity and irreducibility of the semigroup will be invoked repeatedly in constructing positive global solutions and analyzing steady states.

### Lemma 2.2

(Positivity and irreducibility of *e*^𝒜*t*^). *The semigroup* (*e*^𝒜*t*^)_*t*≥0_ *generated by* 𝒜 *on L*^2^(Ω) *is positive and irreducible*.

*Proof*. Consider the bilinear form induced by −𝒜 on the form domain *V* = *H*^1^(Ω). Multiplying −𝒜*n* by *w* ∈ *H*^1^(Ω) and integrating by parts over Ω = (0, 1), with boundary contributions vanishing by the Neumann conditions ∂_*x*_*n*(0) = ∂_*x*_*n*(1) = 0 and the assumption *v*(0) = *v*(1) = 0 from *F*_1_, one obtains

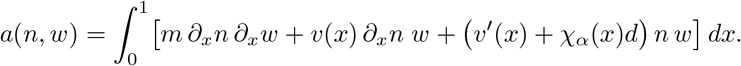

All coefficients appearing in *a*—namely *m >* 0, 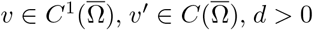, and *χ*_*α*_ ∈ *L*^∞^(Ω)—are real-valued. Since *V* = *H*^1^(Ω) corresponds to Neumann boundary conditions, [11, Proposition 4.4] guarantees the lattice property *n* ∈ *V* ⇒ (Re *n*)^+^ ∈ *V* . These two conditions—real-valued coefficients and the lattice property—are precisely the hypotheses of [11, Theorem 4.2] (see also [11, Corollary 4.3] for the Neumann case), from which it follows that (*e*^𝒜*t*^)_*t*≥0_ is a positive semigroup.

For irreducibility, note that Ω = (0, 1) is connected, 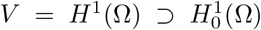, and both the real-coefficient condition and the lattice property have already been verified. These are the hypotheses of [11, Theorem 4.5], which yields the irreducibility of (*e*^𝒜*t*^)_*t*≥0_ in the sense that no closed ideal of *L*^2^(Ω) other than {0} and *L*^2^(Ω) itself is invariant under the semigroup.

We next establish boundedness and positivity of the nonlocal operator 𝒩 defined in (2.2); these properties are essential for the contraction mapping argument in the mild formulation below, and also for the spectral analysis in Section 3.

### Lemma 2.3

(Boundedness and positivity of 𝒩). *The operator* 𝒩 : *L*^2^(Ω) → *L*^2^(Ω) *is a bounded linear operator. Moreover, it is positive in the sense that n* ≥ 0 *implies* 𝒩(*n*) ≥ 0 *a*.*e*..

*Proof*. Linearity is immediate from equation (2.2). To show 𝒩 is bounded, we estimate 𝒩 in two parts. It is easy to see that (1 − *µ*)*χ*_*α*_(*x*)*n*(*x*) is bounded on *L*^2^(Ω), since *χ*_*α*_ ∈ *L*^∞^(Ω) and 0 ≤ 1 − *µ* ≤ 1. Thus

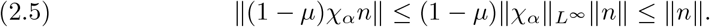

For the integral part, define the operator 𝒦 : *L*^2^(Ω) → *L*^2^(Ω) by

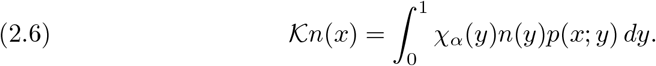

Using Fubini’s theorem and the Cauchy–Schwarz inequality,

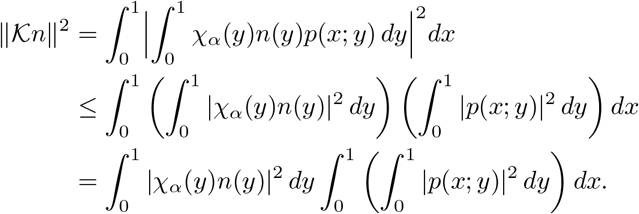

Setting 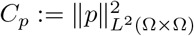, we obtain

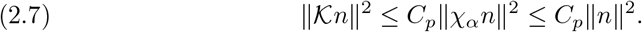

Thus 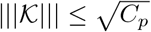. Combining (2.5) and (2.7), we see

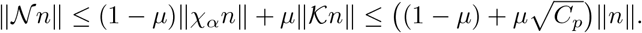

Finally, positivity follows by *n* ≥ 0 with *χ*_*α*_ ≥ 0 and *p* ≥ 0, which implies

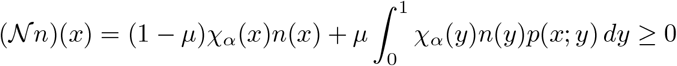

for a.e. *x* ∈ Ω.

With the operator 𝒜 analyzed and 𝒩 shown to be bounded and positive, we are now in a position to rewrite system (1.1)–(1.2) in its mild form and prove that it has a unique local mild solution by using a contraction mapping argument.

### Definition 2.4

(Mild solution). *Let* 0 *< τ < T* . *We say that a pair* (*n, R*) ∈ ℋ_*τ*_ ×𝒫_*τ*_ *is a* mild solution *of* (1.1) *on* [0, *τ*] *with initial data* (*n*_0_, *R*_0_) *if for all t* ∈ [0, *τ*],

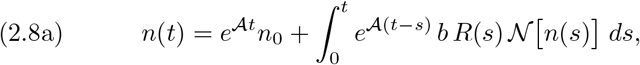

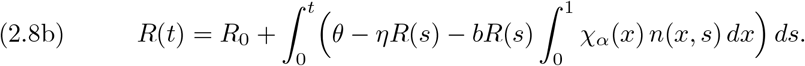

### Theorem 2.5

(Local existence and uniqueness of mild solutions). *Assume that F*_1_*–F*_3_ *hold and that* (*n*_0_, *R*_0_) *satisfy* (1.2). *Then there exists a time τ >* 0 *such that* (1.1) *has a unique mild solution on* [0, *τ*] *with initial data* (*n*_0_, *R*_0_).

*Proof*. For any *τ >* 0, the product space *X*_*τ*_ := ℋ_*τ*_ ×𝒫_*τ*_ with norm 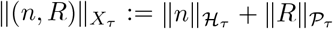 is a Banach space. We define an operator Φ : *X*_*τ*_ → *X*_*τ*_ component-wise by Φ(*n, R*) = (Φ_1_(*n, R*), Φ_2_(*n, R*)), where

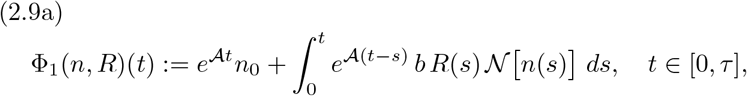

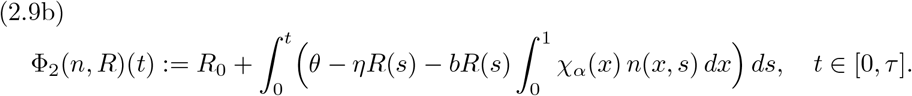

Fixed points of Φ in *X*_*τ*_ are precisely mild solutions of (1.1) on [0, *τ*] with initial data (*n*_0_, *R*_0_). Our goal is to show that for every *r >* 0 there is a time *τ >* 0 such that Φ maps the closed ball of radius *r*

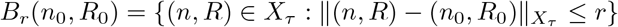

into itself. We then show Φ is a contraction on *B*_*r*_ in order to use the Banach Fixed Point theorem.

Set *M* = ∥*n*_0_∥ + |*R*_0_ | + *r*. Then, we have 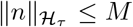 and 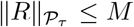 for each (*n, R*) ∈ *B*_*r*_(*n*_0_, *R*_0_). Using the continuity of the semigroup *e*^𝒜*t*^ and Lemma 2.3, we have

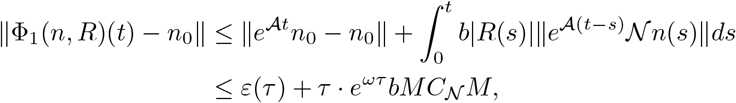

where 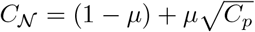 and *ε*(*τ*) → 0 as *τ* → 0. Similarly,

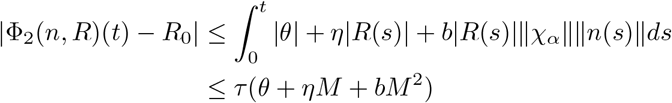

for all *t* ∈ [0, *τ*]. Summing these estimates, we can choose *τ* small enough such that 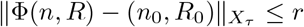. Thus, Φ(*B*_*r*_(*n*_0_, *R*_0_)) ⊆ *B*_*r*_(*n*_0_, *R*_0_).

Next, we prove the map Φ : *B*_*r*_(*n*_0_, *R*_0_) → *B*_*r*_(*n*_0_, *R*_0_) is also a contraction. Let (*n*_1_, *R*_1_), (*n*_2_, *R*_2_) ∈ *B*_*r*_(*n*_0_, *R*_0_). Both pairs are bounded by *M* . Then,

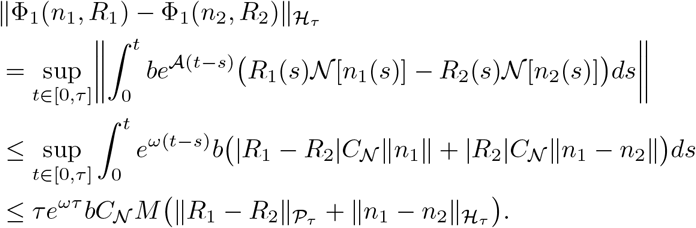

Similarly,

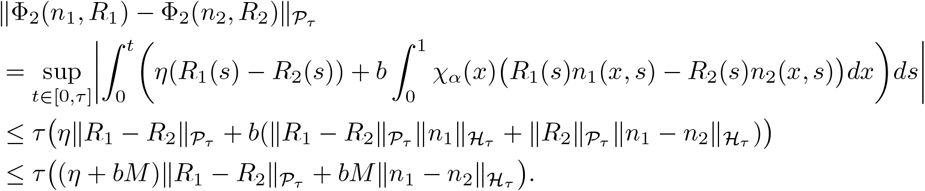

Summing the inequalities gives

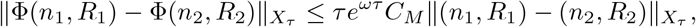

where *C*_*M*_ depends on the constants *M, b, η, C*_𝒩_ . By choosing *τ* such that *τe*^*ωτ*^ *C*_*M*_ *<* 1, the operator Φ is a strict contraction on the complete metric space *B*_*r*_(*n*_0_, *R*_0_). Thus, by the Banach Fixed Point Theorem, there exists a unique fixed point (*n, R*) ∈ *B*_*r*_(*n*_0_, *R*_0_), which is the unique mild solution to system (1.1) on [0, *τ*] with initial data (*n*_0_, *R*_0_).

To extend the result to all *t* ≥ 0, we need the following nonnegativity lemma:

### Lemma 2.6

(Positivity of solutions). *Let τ >* 0 *be the local existence time produced by Theorem 2.5, and let* (*n, R*) *be the unique mild solution on* [0, *τ*] *with initial data* 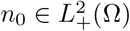 *and R*_0_ ≥ 0 *as in* (1.2). *Then*

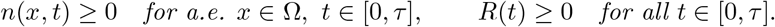

*Proof*. First, define the integrating factor

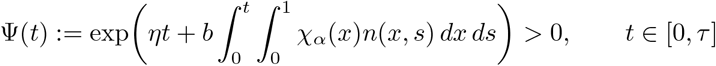

and set *Y* (*t*) = Ψ(*t*)*R*(*t*). Using (1.1b), a direct computation gives, for a.e. *t* ∈ [0, *τ*],

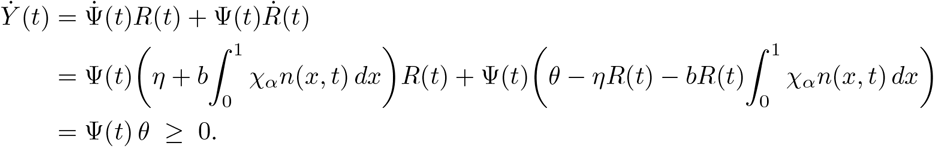

Hence *Y* is nondecreasing on [0, *τ*]. Since *Y* (0) = Ψ(0)*R*_0_ = *R*_0_ ≥ 0, we obtain *Y* (*t*) ≥ 0, and since Ψ(*t*) *>* 0, this gives *R*(*t*) = *Y* (*t*)*/*Ψ(*t*) ≥ 0 for all *t* ∈ [0, *τ*].

Next, define the positive cone 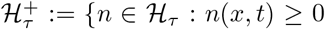 for a.e. *x* ∈ Ω, ∀*t* ∈ [0, *τ*]}. Define a sequence {*n*^(*k*)^(*t*)} by *n*^(0)^(*t*) = *n*_0_ and *n*^(*k*+1)^(*t*) = Φ_1_(*n*^(*k*)^, *R*) where Φ_1_(·, *R*) : ℋ_*τ*_ → ℋ_*τ*_ is defined by (2.9a). By assumption, we have 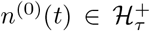. Furthermore, since (*e*^𝒜*t*^)_*t*≥0_ is a positive semigroup by Lemma 2.2, 𝒩 is a positive operator by Lemma 2.3, and *R*(*t*) ≥ 0, then 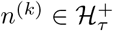 implies

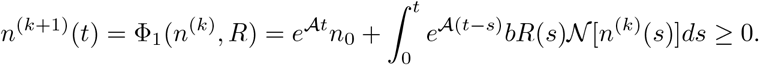

Thus, by induction, 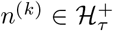 for all *k*. With *R* ∈ 𝒫_*τ*_ fixed, the estimate derived in Theorem 2.5 specializes, taking *R*_1_ = *R*_2_ = *R*, to

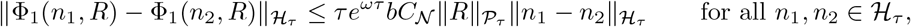

so by the same choice of *τ* that makes Φ a contraction on *B*_*r*_(*n*_0_, *R*_0_), the map Φ_1_(·, *R*) : ℋ_*τ*_ → ℋ_*τ*_ is a strict contraction on ℋ_*τ*_ . Banach’s fixed-point theorem applies on the closed positive cone 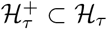, and the iterates 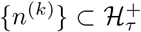 converge to the unique mild solution *n*; since 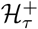 is closed, 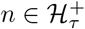.

### Theorem 2.7

(Global well-posedness of mild solutions). *Assume that F*_1_*–F*_3_ *hold and let* (*n*_0_, *R*_0_) *satisfy* (1.2) *with* 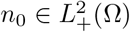 *and R*_0_ ≥ 0. *Then system* (1.1) *has a unique mild solution on* [0, ∞) *and this solution is nonnegative. That is, n*(*x, t*) ≥ 0 *for a*.*e. x* ∈ Ω, *R*(*t*) ≥ 0 *for all t* ≥ 0.

*Proof*. By Theorem 2.5, a unique mild solution (*n, R*) exists on the maximal interval [0, *T*_max_) for some *T*_max_ ∈ (0, ∞]. Lemma 2.6 yields *n* ≥ 0 a.e. and *R* ≥ 0 on the local interval [0, *τ*] produced by Theorem 2.5; applying the same lemma inductively with initial data 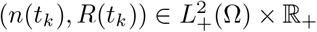 at successive times *t*_*k*_ = *kτ* ∈ [0, *T*_max_) propagates nonnegativity to the entire interval, so *n* ≥ 0 a.e. and *R* ≥ 0 on [0, *T*_max_). We now prove that necessarily *T*_max_ = ∞.

Since *n*(*t*) ≥ 0 for all *t*, we have from (1.1b) that 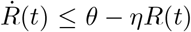 on [0, *T*_max_). By the comparison principle for scalar ODEs,

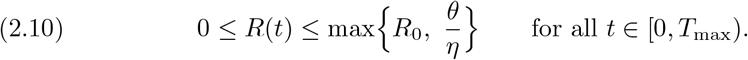

Setting *R*_max_ := max{*R*_0_, *θ/η*}, the mild equation (2.8a) combined with the semigroup estimate (2.3) of Lemma 2.1 and the boundedness of 𝒩 (Lemma 2.3) gives, for *t* ∈ [0, *T*_max_),

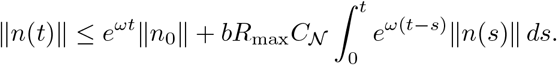

Multiplying by *e*^−*ωt*^ and applying Gronwall’s inequality to *t* ↦ *e*^−*ωt*^∥*n*(*t*)∥ yields

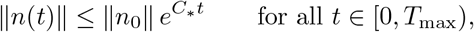

where *C*_∗_ := *ω* + *bR*_max_*C*_𝒩_ depends only on 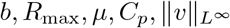, and *m*, but not on *t* or *T*_max_. Combined with (2.10), this gives the uniform a priori bound

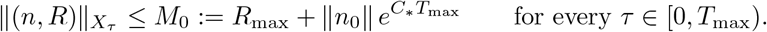

Inspecting the proof of Theorem 2.5, the local existence time is determined by the inequality *τ e*^*ωτ*^ *C*_*M*_ *<* 1, where *C*_*M*_ depends only on a norm bound *M* on (*n, R*) and the model parameters—not on the specific data point. Suppose *T*_max_ *<* ∞. Picking *t*_∗_ *< T*_max_ with *T*_max_ − *t*_∗_ *< τ* (*M*_0_) and applying Theorem 2.5 to the Cauchy problem with data 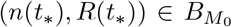 extends the solution strictly beyond *T*_max_, contradicting maximality. Hence *T*_max_ = ∞, and solutions exist for all *t* ∈ [0, ∞).

## 3. Asymptotic Behavior

We now turn our attention to the steady state version of (1.1),

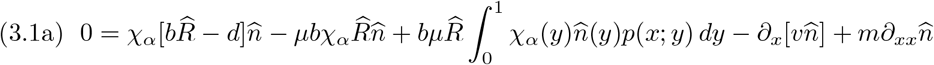

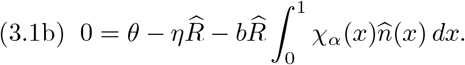

The steady state equation (3.1a) may be written as 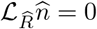, where

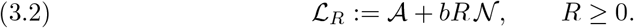

Any nonnegative steady state of (1.1) thus corresponds to a nonnegative element of 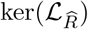 for some 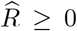. The long-time behavior of solutions is governed by the spectral properties of this family. Our strategy proceeds in several stages: we first establish analyticity and strict positivity for the semigroup generated by 𝒜, then transfer analyticity to ℒ_*R*_ by bounded perturbation and derive compactness of its resolvent. The Krein–Rutman theorem then yields a principal eigenvalue *s*(ℒ_*R*_) for each *R* ≥ 0. We identify the critical level *R* = *d/b* at which *s*(ℒ_*R*_) changes sign by computing the adjoint action on the constant function **1**, and establish strict monotonicity of the map *R* ↦ *s*(ℒ_*R*_). The resulting sign dichotomy determines whether the population goes extinct or persists. We begin with two properties of 𝒜 not required for well-posedness but essential for the spectral analysis.

### Lemma 3.1

(Analytic semigroup generated by 𝒜). *The operator* 𝒜 *generates an analytic C*_0_*-semigroup* {*e*^𝒜*t*^}_*t*≥0_ *on L*^2^(Ω).

*Proof*. The bilinear form associated with − 𝒜 on *V* = *H*^1^(Ω), derived in the proof of Lemma 2.1, is continuous and weakly coercive on *V* . The operator 𝒜 is therefore the operator associated with a continuous, coercive sesquilinear form on the Hilbert triplet *H*^1^(Ω) ⊂ *L*^2^(Ω) ⊂ (*H*^1^(Ω))^∗^, and [19, Theorem 2.18] yields that 𝒜 generates an analytic *C*_0_-semigroup on *L*^2^(Ω).

Combining the positivity and irreducibility established in Lemma 2.2 with the analyticity just proved yields pointwise strict positivity.

### Corollary 3.2

(Strict positivity of *e*^𝒜*t*^). *For every* 0 ≤ *f* ∈ *L*^2^(Ω) *with f* ≢ 0 *and every t >* 0,

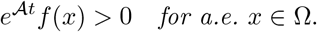

*Proof*. Since (*e*^𝒜*t*^)_*t*≥0_ is positive and irreducible by Lemma 2.2 and holomorphic by Lemma 3.1, the conclusion follows from [1, Theorem 10.1.2].

Since 𝒩 is bounded on *L*^2^(Ω) by Lemma 2.3, the operator *bR*𝒩 is a bounded perturbation of 𝒜 for each fixed *R* ≥ 0. The analyticity established in Lemma 3.1 therefore carries over to the full family (ℒ_*R*_)_*R*≥0_.

### Lemma 3.3

(Analytic semigroup). *For each fixed R* ≥ 0, ℒ_*R*_ *generates an analytic C*_0_*-semigroup on L*^2^(Ω), *and* 𝒟(ℒ_*R*_) = 𝒟(𝒜).

*Proof*. By Lemma 3.1, 𝒜 generates an analytic *C*_0_-semigroup on *L*^2^(Ω), and by Lemma 2.3, 𝒩 ∈ ℒ (*L*^2^(Ω)). Hence *bR*𝒩 is bounded on *L*^2^(Ω), and in particular 𝒜-bounded with 𝒜-bound 0. The perturbation theorem for analytic semigroups [5, Theorem III.2.10] then yields that ℒ_*R*_ = 𝒜+*bR*𝒩 generates an analytic *C*_0_-semigroup on *L*^2^(Ω). Since 𝒩 is defined on all of *L*^2^(Ω), the equality 𝒟 (ℒ_*R*_) = 𝒟 (𝒜) holds by the definition of the sum of operators.

In addition, the spectral analysis below also relies on duality arguments involving the adjoint operators 𝒜^∗^ and 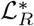, whose domains and actions we now identify.

### Lemma 3.4

(Adjoint of 𝒜 and ℒ_*R*_). *The adjoint of* 𝒜 *is given by*

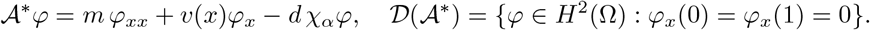

*Moreover, for each fixed* 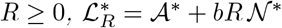, *with* 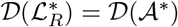, *where*

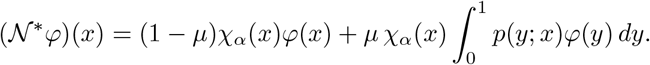

*In particular* 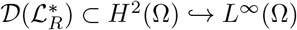

*Proof*. We first identify the adjoint of 𝒩. For *u, φ* ∈ *L*^2^(Ω),

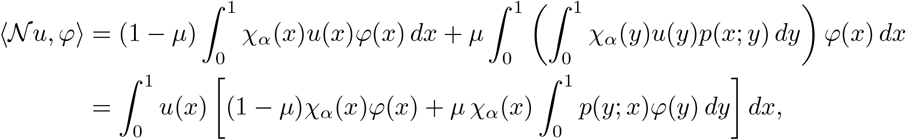

which gives the stated formula for 𝒩 ^∗^*φ*.

We next identify the adjoint of 𝒜. Let *φ* ∈ 𝒟(𝒜^∗^), so there exists *g* ∈ *L*^2^(Ω) with ⟨𝒜*u, φ*⟩ = ⟨*u, g*⟩ for all *u* ∈ 𝒟(𝒜). Taking 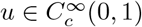, integration by parts yields

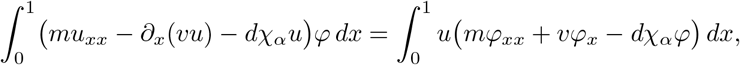

so in the distributional sense *mφ*_*xx*_ + *vφ*_*x*_ − *dχ*_*α*_*φ* = *g* ∈ *L*^2^(Ω). Since *m >* 0, 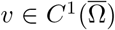, and *χ*_*α*_ ∈ *L*^∞^(Ω), standard one-dimensional elliptic regularity [14, Theorem 8.30] gives *φ* ∈ *H*^2^(Ω).

Now let *u* ∈ 𝒟(𝒜). Integration by parts gives

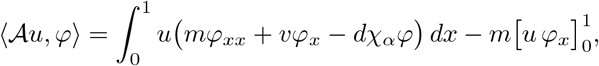

where the transport boundary term vanishes because *v*(0) = *v*(1) = 0. The identity ⟨𝒜*u, φ*⟩ = ⟨*u, g*⟩ then forces the boundary term 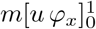 to vanish for all *u* ∈ 𝒟(𝒜). Since *u*(0) and *u*(1) are arbitrary for functions in 𝒟(𝒜), we conclude that *φ*_*x*_(0) = *φ*_*x*_(1) = 0, and therefore

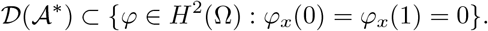

Conversely, if *φ* ∈ *H*^2^(Ω) with *φ*_*x*_(0) = *φ*_*x*_(1) = 0, reversing the above integration by parts shows that

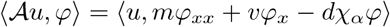

for every *u* ∈ 𝒟(𝒜), so *φ* ∈ 𝒟(𝒜^∗^) with 𝒜^∗^*φ* = *mφ*_*xx*_ + *vφ*_*x*_ − *dχ*_*α*_*φ*. This proves the formula and domain for 𝒜^∗^.

Finally, since *bR*𝒩 is bounded on *L*^2^(Ω), we have 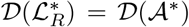 and 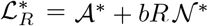, Indeed, if 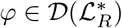, then for all *u* ∈ 𝒟(𝒜),

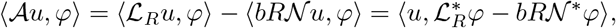

so *φ* ∈ 𝒟 (𝒜 ^∗^), and the reverse inclusion is immediate. The embedding *H*^2^(Ω) → *L*^∞^(Ω) completes the proof.

We can now state the spectral properties of (ℒ_*R*_)_*R*≥0_.

### Proposition 3.5.

*Assume F*_1_*–F*_3_. *Then the following assertions hold for the family* (ℒ_*R*_)_*R*≥0_.

i. *For every R* ≥ 0, *the operator* ℒ_*R*_ *has a compact and positive resolvent*.
ii. *For every R* ≥ 0, *the spectral bound s*(ℒ_*R*_) *is a simple real eigenvalue of* ℒ_*R*_ *admitting a strictly positive eigenfunction. Likewise, s*(ℒ_*R*_) *is an eigenvalue of the adjoint operator* 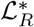 *with a strictly positive eigenfunction*.
iii. *s*(ℒ_*d/b*_) = 0.
iv. *The map R* ↦ *s*(ℒ_*R*_) *is strictly increasing on* [0, *∞*).

*Consequently*,

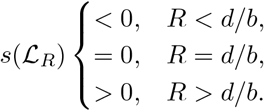

*Proof*. Since 𝒟 (ℒ_*R*_) = 𝒟 (𝒜) (see the proof of Lemma 3.3) and *bR*𝒩 is bounded on *L*^2^(Ω), the graph norms of ℒ_*R*_ and 𝒜 are equivalent. By definition 𝒟 (𝒜) = {*n* ∈ *H*^2^(Ω) : ∂_*x*_*n*(0) = ∂_*x*_*n*(1) = 0 } ⊂ *H*^2^(Ω), so the embedding 𝒟 (𝒜) ↬ *L*^2^(Ω) is compact, and hence so is 𝒟 (ℒ_*R*_) ↬ *L*^2^(Ω). By [5, Proposition II.4.25], ℒ_*R*_ has compact resolvent.

By Lemma 2.2, 𝒜 generates a positive *C*_0_-semigroup on *L*^2^(Ω), and by Lemma 2.3, *bR*𝒩 is a bounded positive operator on *L*^2^(Ω). Hence [5, Corollary VI.1.11] implies that ℒ_*R*_ generates a positive *C*_0_-semigroup 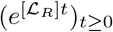 dominating (*e*^𝒜*t*^)_*t*≥0_:

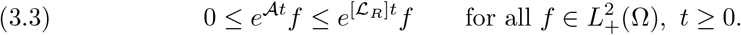

The positivity of the resolvent of ℒ_*R*_ then follows from the Laplace transform representation [5, Theorem II.1.10]

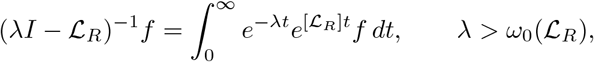

since the integrand is nonnegative for *f* ≥ 0. This proves (i).

For (ii), it suffices to check that the hypotheses of the Krein–Rutman theorem are fulfilled. The irreducibility of 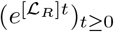 follows from the irreducibility of (*e*^𝒜*t*^)_*t*≥0_ (Lemma 2.2) together with the domination (3.3): for every 0 ≤ *f* ∈ *L*^2^(Ω) with *f* ≢ 0, every *t >* 0, and every measurable *U* ⊂ Ω with |*U* | *>* 0,

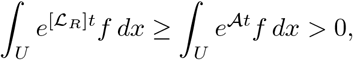

where the strict inequality follows from Corollary 3.2. Hence 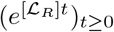 is positive and irreducible.

Since 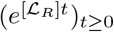 is a positive irreducible *C*_0_-semigroup and ℒ_*R*_ has compact resolvent by (i), the Krein–Rutman theorem [1, Theorem 10.2.5] implies that *s*(ℒ_*R*_) is an eigenvalue of ℒ_*R*_ with a strictly positive eigenfunction *φ*_*R*_ *>* 0, and likewise an eigenvalue of 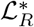 with a strictly positive eigenfunction *ψ*_*R*_ *>* 0. The simplicity of *s*(ℒ_*R*_), with ker(ℒ_*R*_ *− s*(ℒ_*R*_)*I*) = span {*φ*_*R*_ }, follows from [1, Theorem 10.2.6]. This proves (ii).

We now prove (iii). We first identify the action of the adjoint 𝒜^∗^ on the constant function **1** ≡ 1. For every *u* ∈ 𝒟(𝒜),

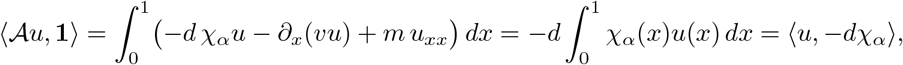

by the assumption *F*_1_ and boundary condition (1.2). Similarly, for the nonlocal operator 𝒩, we have for every *u* ∈ *L*^2^(Ω),

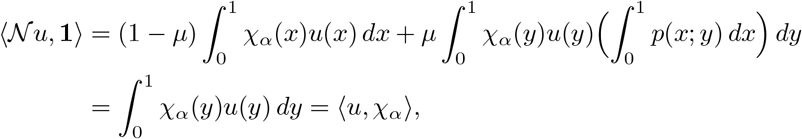

since 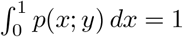 for a.e. *y* ∈ (0, 1) by assumption *F*_2_ . It follows that **1** ∈ 𝒟(𝒜^∗^) with 𝒜^∗^**1** = −*dχ*_*α*_ and 𝒩 ^∗^**1** = *χ*_*α*_. Hence, for *R* = *d/b*,

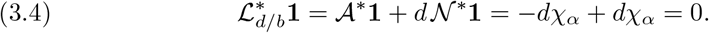

Let *φ*_*d/b*_ *>* 0 satisfy ℒ_*d/b*_*φ*_*d/b*_ = *s*(ℒ_*d/b*_)*φ*_*d/b*_. Pairing with **1** and using (3.4) gives

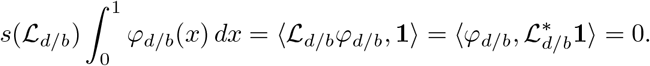

Since *φ* _*d/b*_ *>* 0, 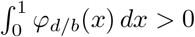, and hence *s*(ℒ _*d/b*_) = 0. This proves (iii).

It remains to prove strict monotonicity of (iv). Let *R*_2_ *> R*_1_ ≥ 0, write 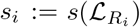, and let *φ*_1_, *ψ*_2_ *>* 0 satisfy

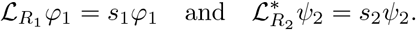

Since 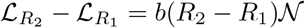, applying this difference to *φ*_1_ gives

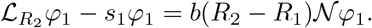

Taking the *L*^2^ inner product with *ψ*_2_ and using

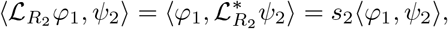

we obtain

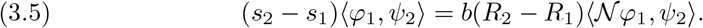

Now ⟨*φ*_1_, *ψ*_2_⟩ *>* 0 since both are strictly positive, and

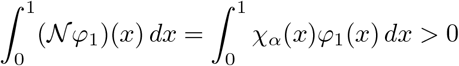

shows 𝒩 *φ*_1_ ≢ 0; therefore ⟨𝒩 *φ*_1_, *ψ*_2_⟩ *>* 0 and (3.5) yields *s*_2_ − *s*_1_ *>* 0. This proves that *R* ↦ *s*(ℒ_*R*_) is strictly increasing. The sign conclusions follow immediately from strict monotonicity together with *s*(ℒ_*d/b*_) = 0.

### Corollary 3.6

(Exponential stability). *Let R*_∗_ *< d/b. Then there exist constants M* ≥ 1 *and ω >* 0 *such that*

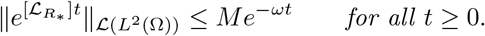

*Proof*. By Proposition 3.5, *R*_∗_ *< d/b* gives 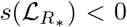. Since 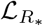 generates an analytic semigroup by Lemma 3.3, its growth bound equals its spectral bound by [5, Corollary IV.3.12]. Therefore the semigroup generated by 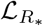 is exponentially stable, and the stated estimate follows (see [5, Proposition VI.1.14 & Theorem VI.1.15]).

The following lemma shows that whenever the active biomass decays to zero, the resource converges to its washout value. This fact is used in the proofs of both washout equilibrium and weak persistence.

### Lemma 3.7

(Resource convergence under active-biomass decay). *Let* (*n, R*) *be a nonnegative mild solution of* (1.1)*–*(1.2). *If*

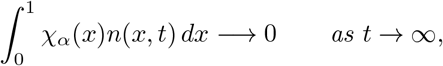

*(then R*(*t*) → *θ/η as t* → ∞.

*Proof*. By the comparison estimate (2.10), *R* is bounded on [0, ∞). Set 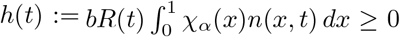, boundedness of *R* and the hypothesis give *h*(*t*) → 0 as *t* → ∞. Variation of constants applied to the scalar equation 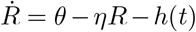 yields

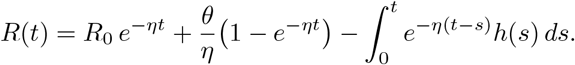

The first two terms tend to 0 and *θ/η* respectively. For the convolution, fix *ε >* 0 and pick *T*_*ε*_ such that *h*(*s*) *< ε* for *s* ≥ *T*_*ε*_; splitting the integral at *T*_*ε*_ and using boundedness 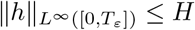,

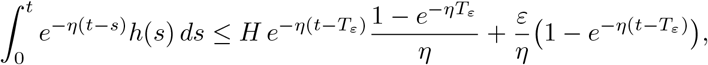

so 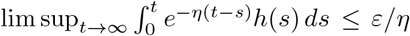 since *ε >* 0 was arbitrary, the convolution tends to 0. Hence *R*(*t*) → *θ/η*.

The asymptotic arguments below involve differentiating projections of the mild solution onto elements of 𝒟 (𝒜^∗^). Since mild solutions need not lie in 𝒟(𝒜), the following lemma provides the required justification.

### Lemma3.8

(Differentiability along adjoint test functions). *Let* (*n, R*) *be a non-negative mild solution of* (1.1)*–*(1.2) *on* [0, *T*). *For every φ* ∈ 𝒟(𝒜^∗^), *the map t* ↦ ⟨*n*(*t*), *φ*⟩ *is continuously differentiable on* (0, *T*) *and satisfies*

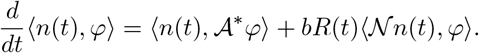

*Proof*. By the mild formula (2.8a) and the duality 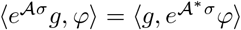,

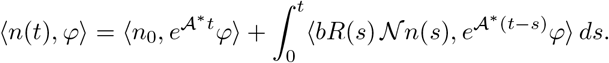

Since *φ* ∈ 𝒟(𝒜^∗^), the map 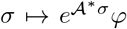 is continuously differentiable in *L*^2^(Ω) with derivative 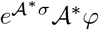, and *s* ↦ *bR*(*s*) 𝒩*n*(*s*) is continuous in *L*^2^(Ω). Differentiating by the Leibniz rule gives

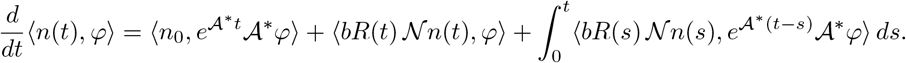

Applying duality to the first and third terms and reassembling via the mild formula (2.8a) yields ⟨*n*(*t*), 𝒜^∗^*φ*⟩ + *bR*(*t*)⟨ 𝒩 *n*(*t*), *φ*⟩, as claimed.

### Theorem 3.9

(Washout equilibrium). *Assume θ/η < d/b. Then* 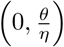 *is the only nonnegative steady state of* (1.1)*–*(1.2). *Moreover, for every nonnegative mild solution* (*n, R*) *of* (1.1)*–*(1.2),

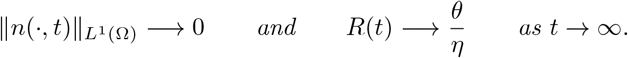

*In particular, the washout state is globally asymptotically stable in the L*^1^(Ω) × ℝ *topology*.

*Proof*. We first show that the washout state is the only nonnegative steady state. Let 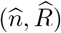 be any nonnegative steady state solution. Integrating (1.1a) over Ω = (0, 1) and using the boundary conditions (1.2), together with *v*(0) = *v*(1) = 0 and 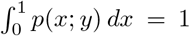, gives 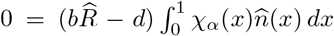. Hence either 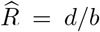 or 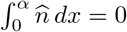. If 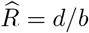, the resource equation yields

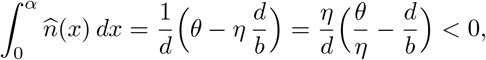

which is impossible since 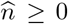. Thus 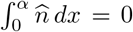, so 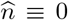 on (0, *α*), and since 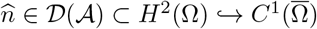, we have 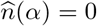. On (*α*, 1) the steady state equation reduces to 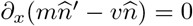, so

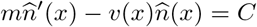

for some constant *C*. Evaluating at *x* = 1 and using 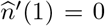 and *v*(1) = 0 gives *C* = 0, and therefore

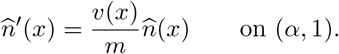

Since *v/m* is continuous on [*α*, 1], uniqueness for this first-order initial-value problem with 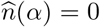 gives 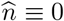 on [*α*, 1]. Thus 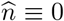 on (0, 1), and the resource equation gives 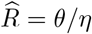. Hence (0, *θ/η*) is the only nonnegative steady state.

We now prove global attraction. Let (*n, R*) be any nonnegative mild solution of (1.1)–(1.2). By the resource equation,

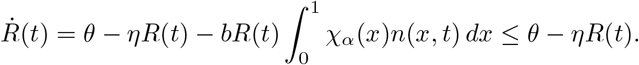

Letting *z* solve *ż* = *θ* − *ηz* with *z*(0) = *R*(0), scalar comparison gives 0 ≤ *R*(*t*) ≤ *z*(*t*) for all *t* ≥ 0. Since *z*(*t*) → *θ/η* and *θ/η < d/b*, we may choose *R*_∗_ with *θ/η < R*_∗_ *< d/b*, so that there exists *T*_0_ *>* 0 with *R*(*t*) ≤ *R*_∗_ for all *t* ≥ *T*_0_.

By Proposition 3.5, the map *R* ↦ *s*(ℒ_*R*_) is strictly increasing with *s*(ℒ_*d/b*_) = 0; since *R*_∗_ *< d/b*, it follows that 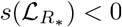, and by Corollary 3.6, there exist constants *M* ≥ 1 and *ω >* 0 such that

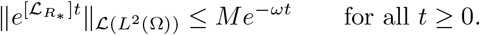

Now fix *t* ≥ *T*_0_. Since *R*(*s*) ≤ *R*_∗_ for all *s* ≥ *T*_0_, we may rewrite the equation (1.1) on [*T*_0_, ∞) as

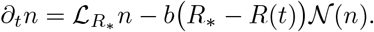

Applying the variation-of-constants formula with generator 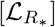 gives

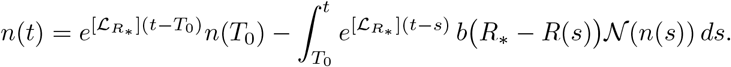

Since *n*(*s*) ≥ 0, 𝒩 is positive, *R*_∗_ − *R*(*s*) ≥ 0, and 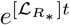 is a positive semigroup, the integral term is nonnegative. Therefore,

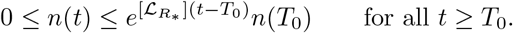

Taking *L*^2^ norms and using the semigroup estimate,

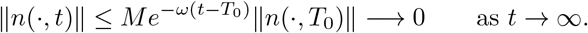

Since |Ω| = 1, we have *L*^2^(Ω) ↬ *L*^1^(Ω) and hence

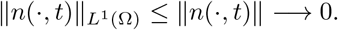

It remains to show that *R*(*t*) → *θ/η*. Since 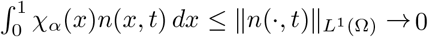, Lemma 3.7 yields *R*(*t*) → *θ/η* as *t* → ∞. Combining with 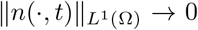, we obtain

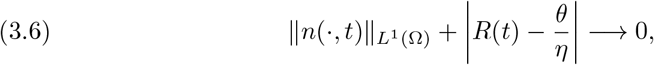

which is the claimed global attraction.

It remains to verify Lyapunov stability in the *L*^1^(Ω) × ℝ topology. With *R*_∗_ as above, set *C* := 1 + *bR*_∗_*/η*. Since **1** ∈ 𝒟 (𝒜^∗^) with 𝒜^∗^**1** = −*d χ*_*α*_ and 𝒩^∗^**1** = *χ*_*α*_ (as shown in the proof of Proposition 3.5(iii)), Lemma 3.8 with *φ* = **1** gives

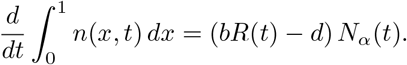

Given *ε >* 0, choose *δ* := min {*ε/*(*C* + 1), *R*_∗_ − *θ/η*} and suppose 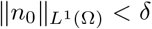 and |*R*_0_ − *θ/η*| *< δ*. The second constraint gives *R*_0_ *< θ/η* + *δ* ≤ *R*_∗_, so (2.10) yields *R*(*t*) ≤ max{*R*_0_, *θ/η*} ≤ *R*_∗_ *< d/b* for all *t* ≥ 0. Hence *bR*(*t*) − *d <* 0, and since *N*_*α*_(*t*) ≥ 0, the total mass is nonincreasing:

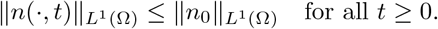

For the resource component, the variation-of-constants formula for 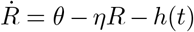 with 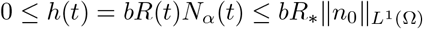 gives

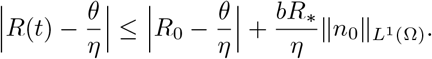

Combining these two estimates,

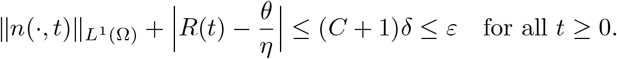

Together with the global attraction established above, this proves global asymptotic stability of the washout equilibrium in *L*^1^(Ω) × ℝ.

### Theorem 3.9

has shown that the system (3.1) converges to the washout equilibrium when *θ/η < d/b*. We now analyze the asymptotic behavior when the opposite inequality, *θ/η > d/b*, holds.

### Theorem 3.10

(Positive equilibrium above threshold). *Assume θ/η > d/b. Then the steady state system* (3.1) *admits exactly two nonnegative equilibria:*

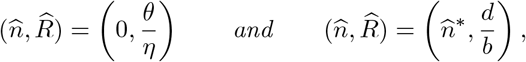

*Where* 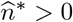.

*Proof*. The washout equilibrium (0, *θ/η*) is an immediate solution of (3.1). Now let 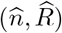 be any nonnegative steady state. Then (3.1a) may be written as 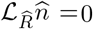. Taking the *L*^2^ inner product of (3.1a) with the constant function **1**, and using 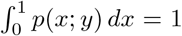, *v*(0) = *v*(1) = 0, and the Neumann boundary conditions (1.2), we obtain

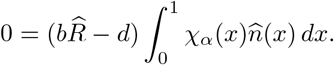

Hence either 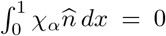 or 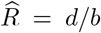 . If 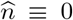, then (3.1b) gives 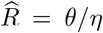, recovering the washout equilibrium.

Assume now that 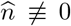. Pairing 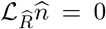 with the strictly positive adjoint eigenfunction 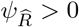 from Proposition 3.5(ii) gives

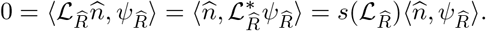

Since 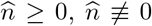, and 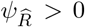 a.e., we have 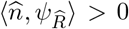, and hence 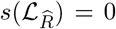. By the strict monotonicity and sign characterization in Proposition 3.5, this forces 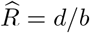. Substituting into (3.1b) gives

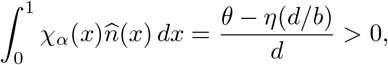

so 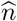 is nontrivial on the active region. At *R* = *d/b*, Proposition 3.5(ii) shows that ker(ℒ_*d/b*_) = span{*φ*_*d/b*_} with *φ*_*d/b*_ *>* 0, so every nonnegative solution of 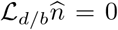 has the form 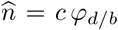 for some *c >* 0. The constant *c* is uniquely determined by (3.1b), namely

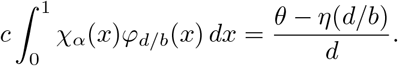

Thus there exists a unique positive equilibrium 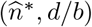, and no other nonnegative steady state is possible.

*Remark* 3.11 (Threshold independence). The critical resource level *R* = *d/b* at which the spectral bound changes sign, and hence the persistence threshold *θ/η* = *d/b*, depend only on the resource and demographic parameters *θ, η, d, b*. In particular, the threshold is independent of the advection velocity *v*, the redistribution kernel *p*, the switching probability *µ*, the persister cutoff *α*, and the diffusion coefficient *m*. Although these parameters shape the internal phenotypic distribution and the profile of the positive equilibrium 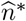, they do not affect the criterion for extinction versus persistence.

Since *s*(ℒ_*θ/η*_) *>* 0 by Proposition 3.5, solutions cannot converge to the washout state when *θ/η > d/b*, which yields the following persistence result.

### Theorem 3.12

(Weak persistence). *Assume θ/η > d/b, and let* (*n, R*) *be a nonnegative mild solution of* (1.1)*–*(1.2) *with n*_0_ ≢ 0. *Then*

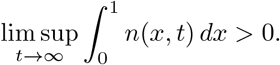

*In particular, n is weakly persistent*.

*Proof*. Set 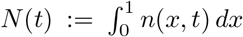 and 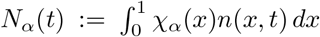. Since 0 ≤ *χ*_*α*_ ≤ 1, we have 0 ≤ *N*_*α*_(*t*) ≤ *N* (*t*) for all *t* ≥ 0. We argue by contradiction: suppose lim sup_*t*→∞_ *N* (*t*) = 0. Then 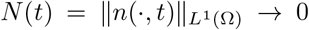 as *t* → ∞, and in particular *N*_*α*_(*t*) → 0; Lemma 3.7 then yields *R*(*t*) → *θ/η*. Choose *R*_−_ with *d/b < R*_−_ *< θ/η*; since *R*(*t*) → *θ/η*, there exists *t*_1_ *>* 0 with *R*(*t*) ≥ *R*_−_ for all *t* ≥ *t*_1_. By Proposition 3.5, 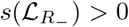, and there exists 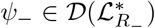 with *ψ*_−_ *>* 0 satisfying

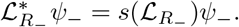

Define *F* (*t*) := *n*(, *t*), *ψ*_−_ . We first justify that *F* (*t*_1_) *>* 0. Since *n*_0_ ≢ 0, the variation-of-constants formula gives

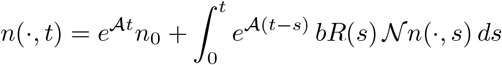

for *t >* 0, and since the integrand is nonnegative, *n*(·, *t*) ≥ *e*^𝒜*t*^*n*_0_. The strict positivity of *e*^𝒜*t*^ (Corollary 3.2) then yields *n*(·, *t*_1_) *>* 0 a.e. on (0, 1), and since *ψ*_−_ *>* 0 a.e., we have *F* (*t*_1_) *>* 0.

Since *ψ*_−_ ∈ 𝒟 (𝒜^∗^) by Lemma 3.4, Lemma 3.8 with *φ* = *ψ*_−_ gives *F* ^*′*^(*t*) = ⟨*n*(·, *t*), 𝒜^∗^*ψ*_−_⟩ +*bR*(*t*)⟨𝒩 *n*(·, *t*), *ψ*_−_⟩. Writing 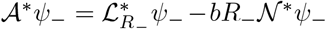 and using the duality ⟨*n*, 𝒩 ^∗^*ψ*_−_⟩ = ⟨𝒩 *n, ψ*_−_⟩, we obtain, for *t* ≥ *t*_1_,

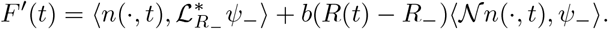

**Fig. 4.1:**
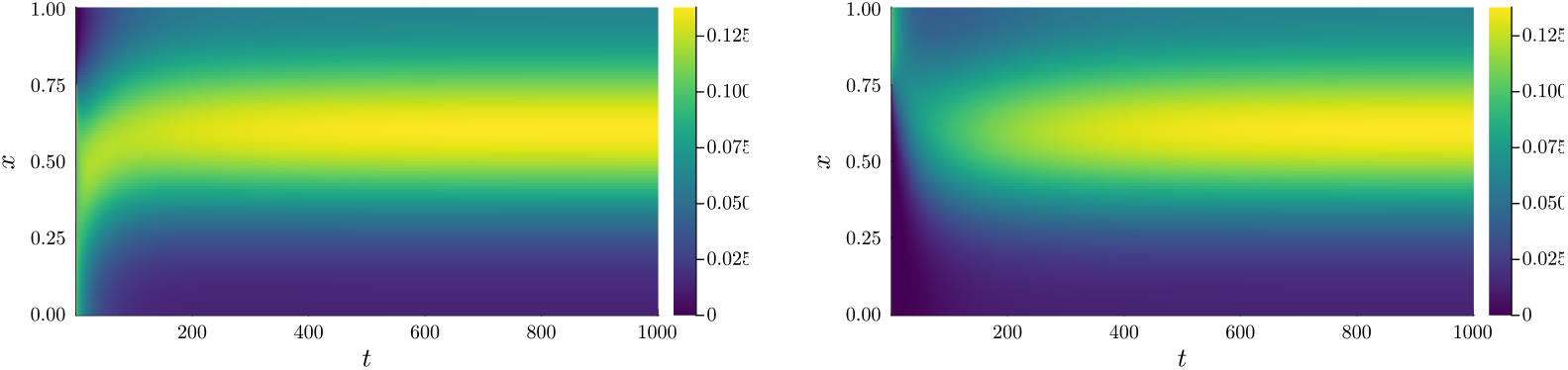
Heatmaps showing the distribution of the persister phenotype *n*(*x, t*) as a function of time under two initial conditions: (a) no persister cells (*n*(*x*, 0) = 0 for *x > α*); (b) only persister cells (*n*(*x*, 0) = 0 for *x < α*). Parameters used were *θ* = 1.0, *η* = 0.3, *d* = 0.01, *b* = 0.6, *µ* = 0.4, *α* = 0.20, *m* = 10^−2^, and drift *v*(*x*) = 60 *x*(*x* − 0.5)(*x* − 1) (homeostatic point *γ* = 0.5).

Since *R*(*t*) ≥ *R*_−_ for *t* ≥ *t*_1_ and both 𝒩 and *ψ*_−_ are positive, the second term is nonnegative, so

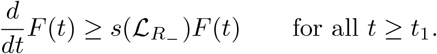

Gronwall’s inequality yields

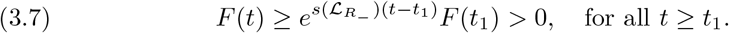

On the other hand, by the Sobolev embedding 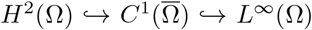 and Lemma 3.4, we have *ψ*_−_ ∈ *L*^∞^(Ω). Hence by Hölder’s inequality,

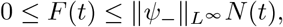

and since *N* (*t*) → 0, we conclude *F* (*t*) → 0, contradicting (3.7).

Hence lim sup 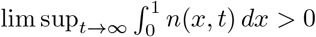, and *n* is weakly persistent.

This result shows that the population is weakly persistent, but the form of that persistence remains unknown. Based on the numerical results in the next section, we have the following conjecture.

### Conjecture 3.13.

*If θ/η > d/b, the unique positive steady state* 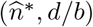 *from Theorem 3.10 is locally asymptotically stable*.

## 4. Numerical Simulations

In this section, we simulate the model by approximating the PDE in equation (1.1) using a system of *K* ordinary differential equations. Code is available at [Tyler Meadows’s GitHub repository](https://github.com/Tyler-Meadows/Persistence).

For simplicity, we assume that *α* = *k/K* for some *k, K* ∈ 𝒩 with *k < K*, so that no compartment contains both persister and regular cells. Each function may be approximated by its value at the midpoint of the compartment, denoted *x*_*i*_ for the *i*th compartment. Spatial derivatives are approximated by central finite differences,

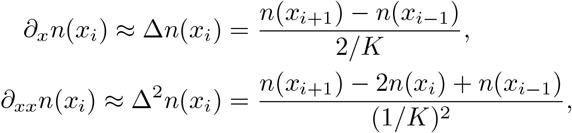

with the Neumann boundary conditions imposed through ghost cells *n*_0_ := *n*_1_ and *n*_*K*+1_ := *n*_*K*_. The integral term is approximated by a sum,

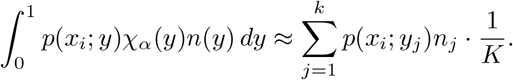

Writing *n*(*x*_*i*_, *t*) = *n*_*i*_, the finite-difference approximation of (1.1) reads

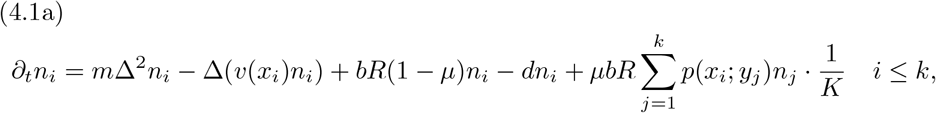

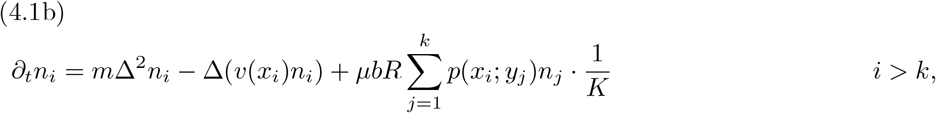

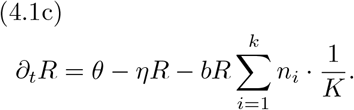

In our simulations we take *p*(*x*; *y*) ≡ 1, so that daughter cells are uniformly distributed over all phenotypes. Figure 4.1 shows convergence to the positive steady state under two initial conditions: (a) no persister cells; (b) only persister cells. Time series simulations were done using the differential equations package in Julia [13]. These numerical results are consistent with Conjecture 3.13; convergence from two markedly different initial conditions to the same steady state further suggests that global asymptotic stability may hold, a stronger conjecture we leave for future work.

## Acknowledgments

The authors thank Adrian Lam and Thomas Hillen for many fruitful discussions. T. Day is supported by an NSERC Discovery Grant.

